# Population structure and pharmacogenomic risk stratification in the United States

**DOI:** 10.1101/2020.03.28.013755

**Authors:** Shashwat Deepali Nagar, Andrew B. Conley, I. King Jordan

## Abstract

Pharmacogenomic (PGx) variants mediate how individuals respond to medication, and response differences among racial/ethnic groups have been attributed to patterns of PGx diversity. We hypothesized that genetic ancestry (GA) would provide higher resolution for stratifying PGx risk, since it serves as a more reliable surrogate for genetic diversity than self-identified race/ethnicity (SIRE), which includes a substantial social component. We analyzed a cohort of 8,628 individuals from the United States (US), for whom we had both SIRE information and whole genome genotypes, with a focus on the three largest SIRE groups in the US: White, Black, and Hispanic. Whole genome genotypes were used to characterize individuals’ continental ancestry fractions – European, African, and Native American – and individuals were grouped according to their GA profiles. SIRE and GA groups were found to be highly concordant. Continental ancestry predicts individuals’ SIRE with >96% accuracy, and accordingly GA provides only a marginal increase in resolution for PGx risk stratification. PGx variants are highly diverged compared to the genomic background; 82 variants show significant frequency differences among SIRE groups, and genome-wide patterns of PGx variation are almost entirely concordant with SIRE. Nevertheless, 97% of PGx variation is found within rather than between groups. Examples of highly differentiated PGx variants illustrate how SIRE partitions PGx variation based on group-specific ancestry patterns and contains valuable information for risk stratification. Finally, we show that individuals who identify as Black or Hispanic benefit more when SIRE is considered for treatment decisions than individuals from the majority White population.

## Introduction

Pharmacogenomic (PGx) variants are associated with inter-individual differences in drug exposure and response, affecting medication dosage, efficacy and toxicity^1; 2^. A number of studies have shown racial and/or ethnic differences in drug response^3–7^, based in part on group-specific differences in the frequencies of PGx variants^8^. A 2015 review found that 20% of drugs approved over the previous six years showed response differences among racial/ethnic groups, and these differences are often translated into group-specific prescription recommendations that are issued on FDA-approved drug labels^7^. Examples of such recommendations include contraindication of Rasburicase, a medication used to clear uric acid from the blood in patients undergoing chemotherapy, for individuals of African or Mediterranean ancestry, and a toxicity warning for the anticonvulsant Carbamazepine in Asian patients. A higher dosage of the immunosuppressive drug Tacrolimus is indicated for African-American transplant patients, whereas a lower initial dose of Rosuvastatin is recommended for Asians. Despite the inclusion group-specific recommendations in a number of drug labels, the utility of racial and ethnic categories in biomedical research, and their relevance to clinical decision making, remain a matter of substantial controversy^9–12^.

Critiques of the use of racial and ethnic categories in biomedical research point to the appalling history of race science^13–15^ and stress the potential of such research to reify outmoded notions of racial difference^16–18^. This school of thought holds that race is a primarily a social construct with little or no biological (genetic) meaning^19–23^. As it relates to clinically relevant PGx variation across groups, the extent to which racial and ethnic categories serve as a reliable proxy for genetic diversity has also been called into question. The authors of the recent commentary ‘Taking race out of human genetics’ make a compelling case for eliminating the use of race as a category in genetic research, asserting that race and ethnicity are taxonomic (*i.e.* categorical) labels that by definition cannot capture the full complexity of individuals’ genetic ancestry^24^. They suggest that genetics research should instead focus on biogeographically defined populations and genetic ancestry, as opposed to racial categories, and for this study we hypothesized that genetic ancestry should better partition PGx variation than SIRE. We posit that genetic ancestry provides a number of advantages over racial/ethnic categories for biomedical research: (i) it can be characterized independent of the social and environmental dimensions of race/ethnicity, (ii) it can be measured objectively and with precision, and (iii) it can be quantified as a continuous variable, as opposed to categorical racial/ethnic labels. Indeed, a number of recent studies have focused on PGx variation among populations defined by genetic ancestry rather than racial and ethnic groups^25–30^.

The goal of this study was to compare the relative utility of race/ethnicity versus genetic ancestry for partitioning PGx variation among populations in the United States (US). We focused on individuals aged 50 and older, 75% of whom take prescription medication on a regular basis^31^, and restricted our study to the three largest racial/ethnic groups in the US: White, Black (or African-American), and Hispanic/Latino^32^. Our study cohort is made up of 8,629 participants from the Health and Retirement Study (HRS)^33^, for whom we had both SIRE information and whole genome genotypes. We first compared the relationship between self-identified race/ethnicity (SIRE) and genetic ancestry (GA), characterized via analysis of whole genome genotype data, and we then measured the extent to which PGx variation is partitioned by SIRE versus GA. We provide a number of examples of PGx variants that are highly differentiated among groups and discuss the implications of these findings in light of population genetics and clinical decision-making.

## Materials and Methods

### Study Cohort

Self-identified race and ethnicity (SIRE) information and whole genome genotypes for Americans over the age of 50 and their spouses were collected as part of a nationally-representative longitudinal panel study called the Health and Retirement Study (HRS)^33^. For the current study, only HRS participants with both SIRE and genotype information were considered (8,912 participants). The 284 participants who did not identify with one of the three largest racial/ethnic categories in the HRS data – non-Hispanic White (5,927), non-Hispanic Black (1,527), and Hispanic/Latino of any race (1,174) – were excluded from this analysis. This yielded a total of 8,628 individuals in our final analysis cohort.

### Genetic Ancestry (GA) Analysis

HRS participants were previously genotyped at ~2,381,000 genomic sites using the Illumina Omni2.5 BeadChip^33^. Whole genome genotype data from HRS participants were compared to reference populations from Europe, Africa, and the Americas in order to infer their continental genetic ancestry patterns as previously described (Supplementary Table 1)^34^. Reference populations were taken from (i) the 1000 Genomes Project (648)^35^, (ii) the Human Genome Diversity Project (110)^36^, and (iii) 21 Native American populations from across the Americas (90)^37^. A custom script that employs PLINK version 1.9^38^ was used to harmonize the HRS and reference population variant calls. The variant call data were merged by identifying the set of variants common to both datasets, with strand flips and variant identifier inconsistencies corrected as needed. The initial merged and cleaned variant data set was filtered for variants with >1% missingness and <1% minor allele frequency among samples. The final harmonized genotype data contains 228,190 genomic sites. The harmonized genotype dataset was phased using ShapeIT version 2.r837^39^. ShapeIT was run without reference haplotypes, and all individuals were phased at the same time. Individual chromosomes were phased separately, and the X chromosome was phased with the additional ‘-X’ flag.

A modified version of the RFMix program^34^;^40^ was used to characterize the continental genetic ancestry patterns for the HRS participants, with European, African, and Native American populations used as reference populations. RFMix was run in the ‘PopPhased’ mode with a minimum node size of five, using 12 generations and the “—use-reference-panels-in-EM” for two rounds of EM, to assign continental ancestry for haplotypes genome-wide. Contiguous regions of ancestral assignment, “ancestry tracts,” were created where RFMix ancestral certainty was at least 95%, and genome-wide continental ancestry estimates for HRS participants were obtained by averaging across confidently assigned ancestry tracts.

Non-overlapping genetic ancestry (GA) groups were defined from individual participants’ continental ancestry estimates obtained via RFMix analysis using *k-* means clustering implemented in the Python package Scikit-learn^41^ with *k*=3. Each participant was represented as a point in three-dimensional (3-D) space, parameterized by their three continental ancestry fractions. Formally, the position of a participant (*i*) in this genetic ancestry space was defined by (*E_i_*, *A_i_*, *N_i_*), where *E_i_*, *A_i_*, and *N_i_* are the European, African, and Native American ancestry fractions. *K*-means clustering using Euclidean distances between all pairs of individual participants in this 3-D genetic ancestry space to yield three non-overlapping clusters. Given that *k*-means clustering can be unstable, the algorithm was run on these data 100 times and the most probable group membership was assigned to each participant. This method allowed us to define three non-overlapping groups of HRS participants informed entirely by their genetic ancestry and free from the social dimensions of SIRE.

The association between GA and PGx variant genotypes was measured using our previously described method^25^. To obtain the strength of association (*β*) between continental ancestry proportions and genotypes, continental ancestry fractions were regressed against the observed PGx variant genotypes. Formally, the genetic ancestry fraction *y* = *βx* + *ε*, where *x* ∈ {0,1,2} refers to the number of PGx variant effect alleles. The significance of these ancestry associations was quantified using a t-test.

### Measurement of PGx Variation

Single nucleotide variants (SNVs) associated with pharmacogenomic response – *i.e.* PGx variants – were mined from the Pharmacogenomic Knowledgebase (PharmGKB)^2^. This online database is a source of manually curated clinical variant annotations for PGx variants and their associated drug-response phenotypes. Data on the chromosomal locations of PGx variants, the identity of PGx effect (risk) alleles, PGx variants’ mode of effect (additive or dominant), clinical annotations, and clinical evidence levels were parsed and taken for analysis. A total of 2,351 PGx variants were accessed from PharmGKB, 989 of which were genotyped for the HRS cohort. PharmGKB annotates the specific effect alleles that are associated with interindividual differences in drug dosage, efficacy, and toxicity. The direction of effect (higher or lower) is specific to individual PGx variants for dosage and efficacy, whereas toxicity effect alleles always correspond to increased toxicity.

PGx allele frequencies for SIRE and GA groups were computed as the group-specific counts of effect alleles normalized by the total number of typed individuals for each group. Pairwise between group fixation index (*F_ST_*) values for each variant were computed by calculating two components: (i) the mean expected heterozygosity within subpopulations, 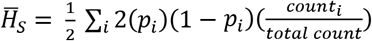, where *p_i_* is the frequency of risk allele in population *i*, and *count_i_* is the number of individuals in population *i*, and *total count* refers to the total number of individuals in both populations and (ii) the expected heterozygosity in the total population, 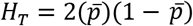, where 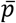 is the mean effect allele frequency in both populations under consideration. The fixation index was computed by combining the two computed metrics as 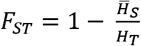 ^42^. PGx variants were used to calculate pairwise inter-individual distances for all HRS participants using PLINK, and the resulting distance matrix was projected into two dimensions using multi-dimensional scaling (MDS) with the mds function in R. *K*-means clustering of the participants in MDS space was used to generate three non-overlapping PGx variant groups in the same way as described for the GA groups.

Odds ratios (*ORs*) were calculated for group-specific PGx effect allele counts^43^. In a contingency table for the counts of effect allele in population P_A_ with the four values: P_E_ (Effect allele count in P_A_), P_N_ (Non-effect allele count in P_A_), Q_E_ (Effect allele count in non-PA individuals), Q_N_ (Non-effect allele count in non-P_A_ individuals), this was done using the formula 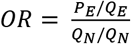, with confidence intervals calculated as *CI* = exp(*log(OR*) ± *Z_α/2_* * *SE_log(oR)_*), where *α* is 0.05, *Z_α/2_* is 1.6, and 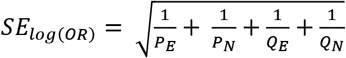. Similarly, using group-specific PGx effect counts the absolute risk increase (*ARI*) was calculated as *ARI* = 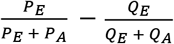, with confidence intervals calculated as *CI* = *ARI* ± *Z_α/2_ × *SE*_ARI_*, where α is 0.05, *Z_α>/2_* is 1.96, and 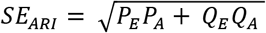^44^. Group-specific genotype prediction accuracy values were calculated as *Accuracy* = (*TP* + *TN*)/(*TP* + *TN* + *FP* + *FN*), where *TP* is true positives, *TN* is true negatives, *FP* is false positives, and *FN* is false negatives. *TP, TN, FP*, and *FN* designations are assigned based on the SIRE group that shows enrichment for PGx effect allele (or genotype). The presence of the PGx effect allele in the implicated SIRE group is counted as a true positive, whereas its presence in the other groups is counted as a false positive. Conversely, the presence of the PGx non-effect allele in the implicated SIRE group is counted as a false negative, whereas its presence in the other groups is counted as a true negative. Accuracy confidence intervals are calculated as 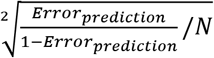, where 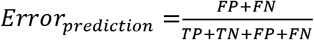 and *N* × *TP* + *TN* + *FP* + *FN*. As noted before, when α is 0.05, *Z_α/2_* is 1.96.

Pre- and post-test probabilities were compared in order to compute the amount of information gained per 100 individuals based on PGx stratification with SIRE. For any given PGx variant, the pre-test probability is calculated as the overall population prevalence of the PGx effect allele (additive mode) or genotype (dominant mode): *Prevalence_overall_* = *Count_EA_/Count_Total_*, where *Count_EA_* is the count of the effect allele/genotype in the cohort and *Count_Total_* is the total count of alleles/genotypes at that locus in the cohort. The post-test probability is calculated as the group-specific positive predictive values (PPVs) for the PGx effect allele or genotype. *PPV* is calculate as: 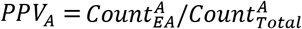, where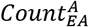 is the count of the effect allele/genotype in population *A* and 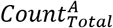 is the total count of alleles/genotypes at that locus in the population *A*. Information gain is then calculated as: *lnfoGain_A_* = *|PPV_A_* — *Prevalence_overall_|*.

### Comparison of SIRE and GA

To test whether PGx variant allele frequencies were correlated between SIRE and GA, pairwise PGx variant allele frequency differences calculated for SIRE groups were regressed against allele frequency differences calculated for GA groups. Here, the null hypothesis is *H_0_: β=* 0, while the alternate hypothesis is *H_A_: β≠* 0. The significance of this correlation was testing using a t-test where *t* = (*β_obs_* – *β_exp_*)/*SE* and *P* = *P*(*T_DF_* ≤ *β_exp_*). Next, we tested whether GA groups partition PGx variation more than SIRE groups using the same regression. For this test, the null hypothesis is *H_0_*: β = 1, while the alternate hypothesis is *H_A_*: β < 0. An underlying assumption for this one-tailed test is that GA groups should hold more information about PGx allele frequency differences when compared to SIRE groups. We calculated the difference in the expected (unity line) and observed (SIRE versus GA) regression slopes, *d* = (*β_exp_* – *β_obs_*)/2 to quantify the magnitude of the effect. A denominator of 2 was chosen to reflect the entire range of possible slopes that the data may take – going from –1, where SIRE groups reflect exactly the opposite difference in allele frequencies, to 1, where SIRE groups faithfully and completely capture the allele frequency differences observed in GA groups. The statistical significance was tested using a t-test as described above.

## Results

### Self-identified race/ethnicity (SIRE) and Genetic Ancestry (GA) in the US

We compared SIRE to GA for a cohort of 8,628 individuals characterized as part of the Health and Retirement Study (HRS), for whom both SIRE information and whole genome genotypes were available (Table 1). HRS participants self-identified according to racial and ethnic labels defined by the US Government Office of Management and Budget (OMB). OMB defines five racial groups and two ethnic groups to assess disparities in health and environmental risks^45^. HRS participants were asked to select one or more race category and a single ethnic designation as Hispanic/Latino or not. We considered the race and ethnicity selections together and focused on the three largest categories in the HRS cohort: non-Hispanic White (5,927; 68.7%), non-Hispanic Black (1,527; 17.7%), and Hispanic/Latino of any race (1,174; 13.6%). We refer to these three groups here as White, Black, and Hispanic. The percentages of each SIRE group in the HRS cohort resemble the demographics of the US: White=72.4%, Black=12.6%, and Hispanic=16.3%^45^.

**Table 1.**
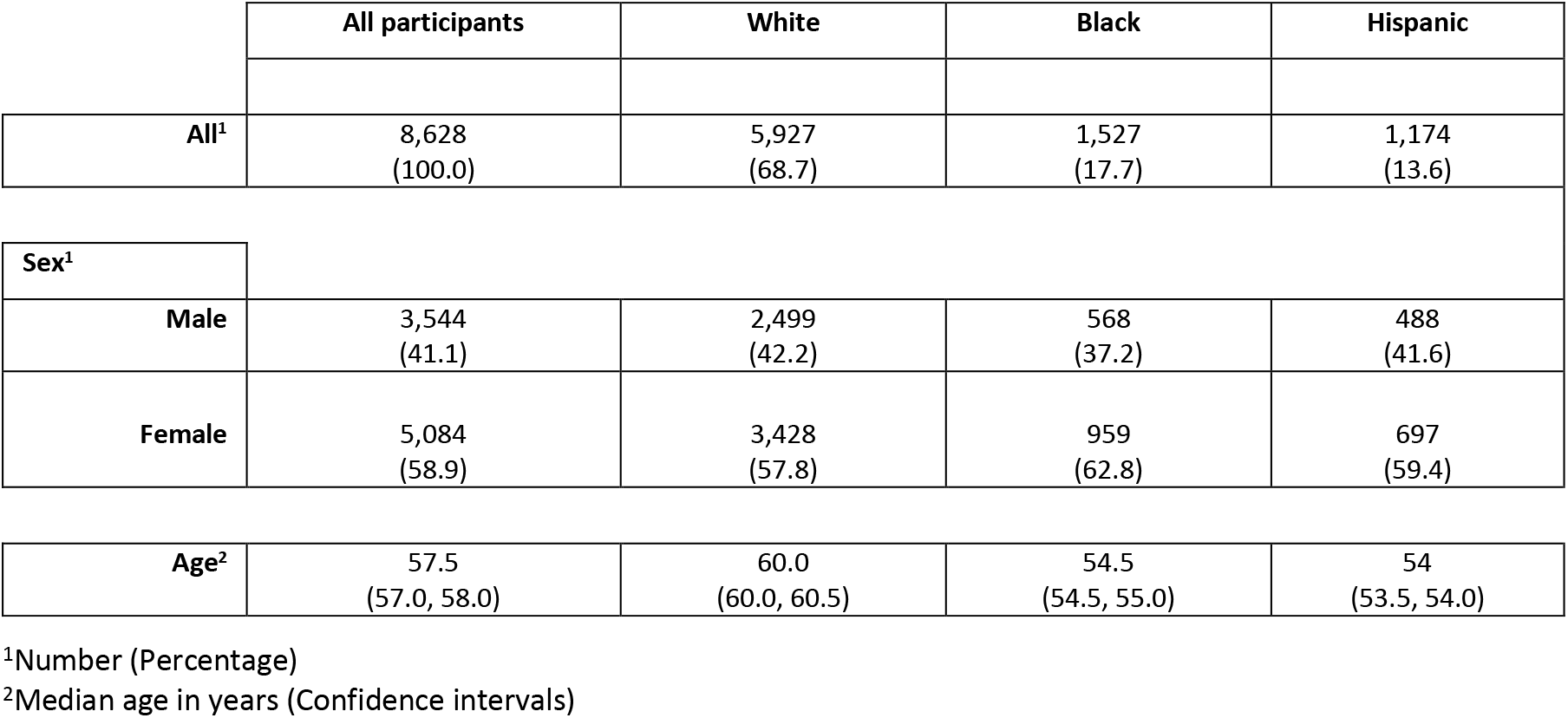
Demographic description for the cohort used in this study.

Continental ancestry profiles were inferred for members of the HRS cohort by comparing their whole genome genotypes to whole genome sequence and genotype data for reference populations from Europe, Africa, and the Americas as described in the Materials and Methods. Each HRS participant was assigned European, African, and Native American ancestry proportions, and the resulting ancestry profiles were then clustered into three distinct (non-overlapping) GA groups using *k*-means clustering. GA groups were defined without reference to SIRE group labels, using unsupervised clustering on continental ancestry fractions alone, and the choice to cluster ancestry profiles into three groups was made to allow for direct comparison with the three SIRE groups and in light of known patterns of continental ancestry in the US^46^. Permutation analysis was used to confirm the stability of the resulting GA groups and their robustness to changes in sample size (Supplementary Figure 1). The distributions of continental ancestry fractions were compared for the three SIRE groups – White, Black, and Hispanic – and the three GA groups (Figure 1).

**Figure 1.**
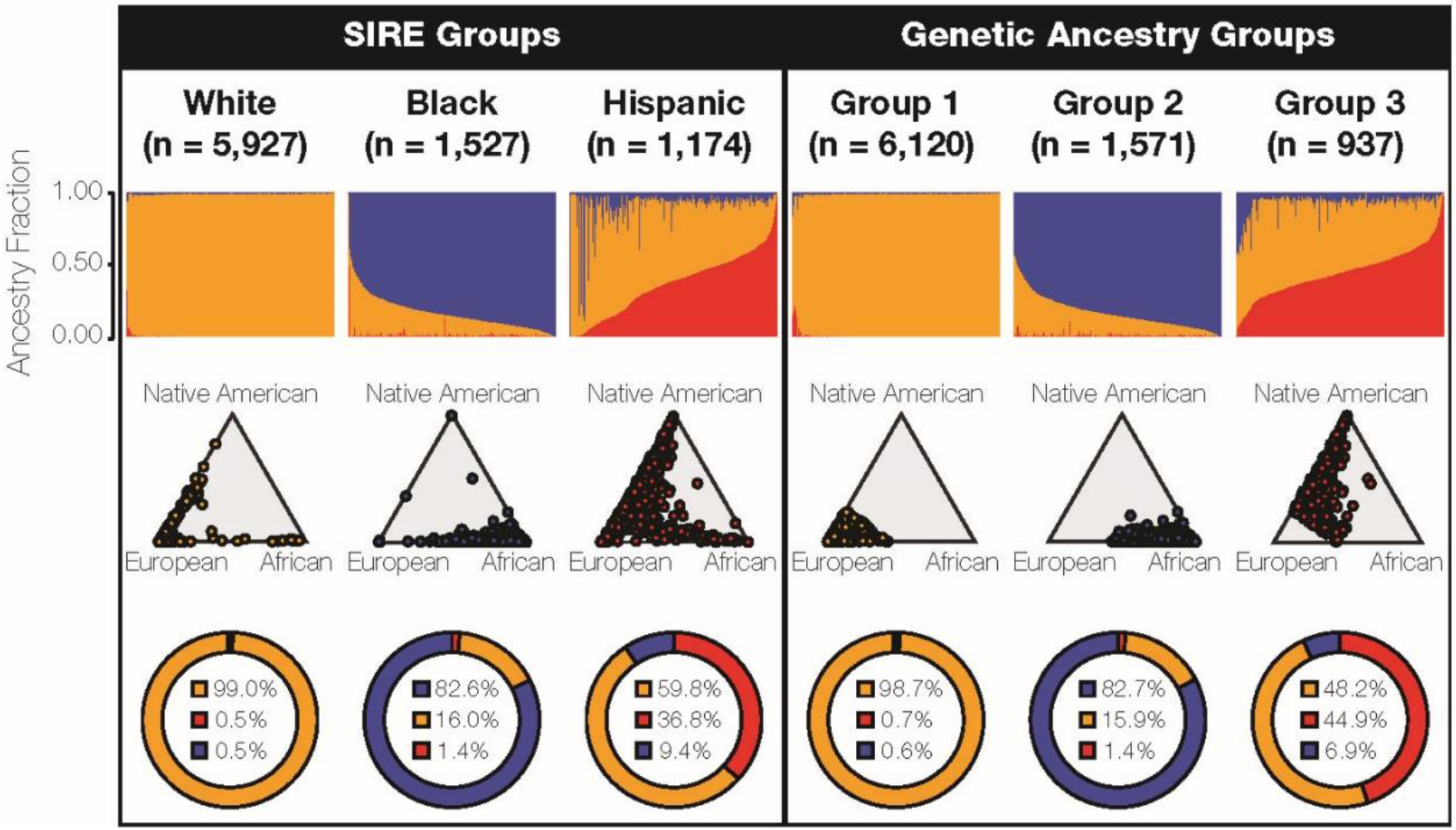
Race, ethnicity, and genetic ancestry in the US. Continental genetic ancestry patterns are shown for self-identified race/ethnicity (SIRE) and genetic ancestry (GA) groups: European ancestry (orange), African ancestry (blue), and Native American ancestry (red). HRS cohort participants are grouped by SIRE and GA, as described in the text, and continental ancestry fractions are compared for each grouping system. Top row: continental ancestry fractions for individuals organized into the three SIRE and three GA groups. Each column represents an individual genome, and the three continental ancestry fractions are shown for each individual column. Middle row: ternary plots showing the continental ancestry fractions for the SIRE and GA groups, as illustrated by the relative proximity to each of the three ancestry poles. Bottom row: average continental ancestry percentages for the SIRE and GA groups.

**Figure 2.**
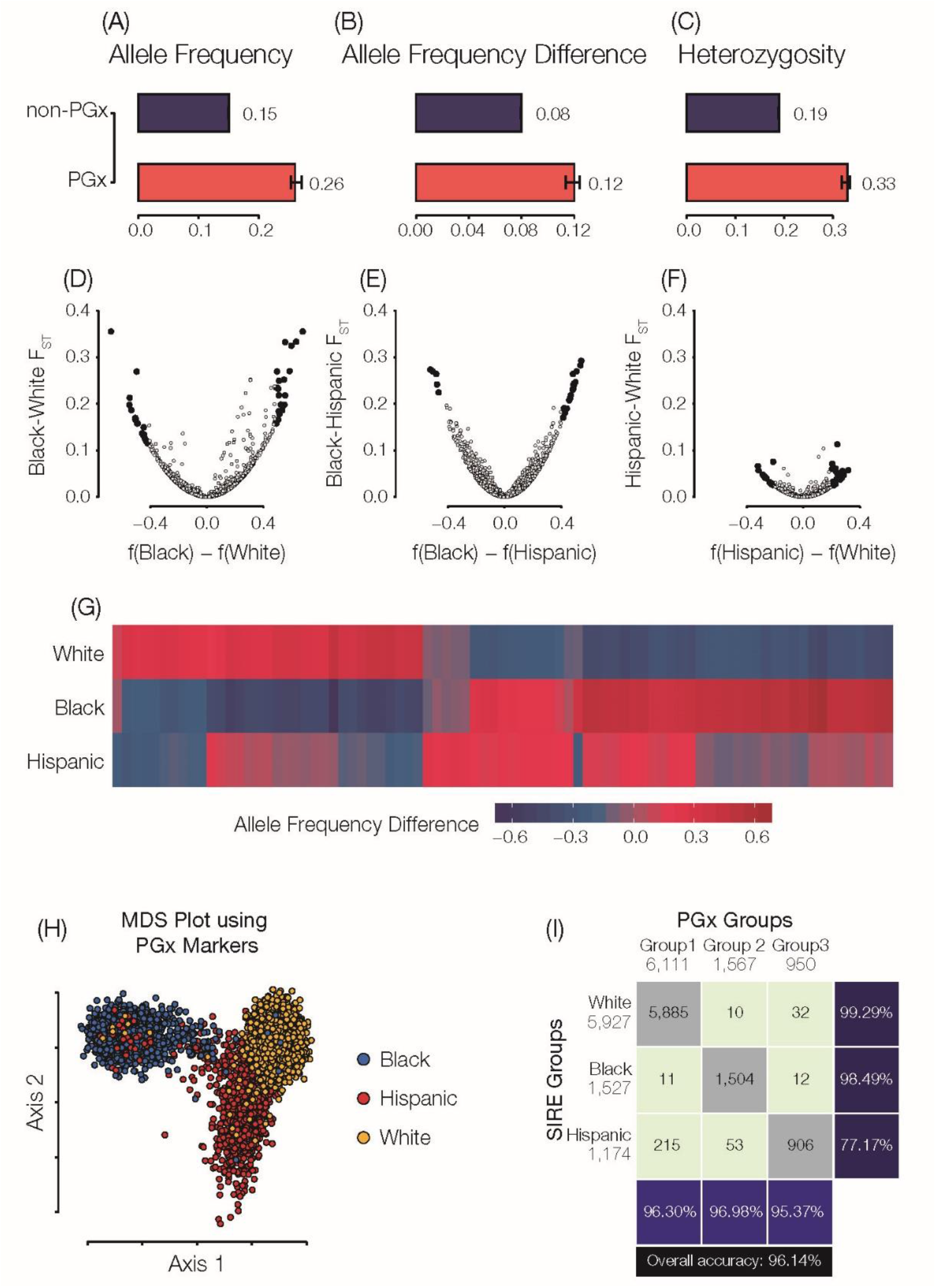
Pharmacogenomic variation in the US. Genome-wide average allele frequencies (A), group-specific allele frequency differences (B), and heterozygosity fractions (C) are shown for PGx variants (red) compared to non-PGx variants (blue). (D-F) Fixation index (FST; y-axis) and allele frequency differences (x-axis) for pairs of SIRE groups. Statistically significant PGx allele frequency differences are highlighted in black. (G) Heatmap showing group-specific allele frequencies for significantly diverged PGx variants. (H) Multi-dimensional scaling (MDS) plot showing the relationship among individual genomes as measured by PGx variants alone. Each dot is an individual HRS participant genome, and genomes are color-coded by participants SIRE. (I) The correspondence between SIRE groups and PGx groups defined by K-means clustering on the results of the MDS analysis. Data shown here correspond to SIRE groups; analogous results for GA groups are shown in Supplementary Figure 4.

The three objectively defined GA groups appear to correspond well to the SIRE groups, with respect to the distributions of individuals’ continental ancestry fractions (Figure 1 – top row). GA groups 1, 2, and 3 correspond to the White, Black, and Hispanic SIRE groups, respectively. The distributions of continental ancestry fractions for the SIRE and their corresponding GA groups are compared in Supplementary Figure 2. Despite the apparent similarity between SIRE and GA, ternary plots underscore the broader distribution of ancestry fractions within SIRE groups compared to the non-overlapping GA groups delineated by *k*-means clustering (Figure 1 – middle row). This is especially true for the Hispanic group, consistent with the fact that it may include individuals who identify as any race. Overall, SIRE and the GA groups show similar average continental ancestry percentages: White/Group 1 show ~99% European ancestry, Black/Group2 have ~82% African ancestry, and Hispanic/Group 3 show predominantly European ancestry (~60%) with the highest levels of Native American ancestry (~37%) and the greatest variance in continental ancestry for any of the three groups.

The correspondence between the SIRE and GA groups was quantified by characterizing the overlap of membership assignments across the two groupings (Supplementary Figure 3). Overall, individuals’ membership in the three SIRE and corresponding GA groups show 96.2% concordance. The highest concordance is seen for the White/Group 1 pair, followed by Black/Group 2, with Hispanic/Group 3 showing the lowest concordance. The levels of concordance vary according to which grouping system is taken as the reference for comparison. This distinction is most obvious for the Hispanic/Group 3 pairing: 96.6% of Group 3 members self-identify as Hispanic, while only 77.1% of self-identified Hispanics fall into Group 3.

### Pharmacogenomic variation in the US

PGx variants that influence drug response were mined from the PharmGKB database, and levels of PGx variation were compared within and between the SIRE and GA groups defined for the HRS cohort. Results for SIRE group comparisons are shown in Figure 2, and results for the analogous comparison of GA groups are shown in Supplementary Figure 4. PGx variants show higher allele frequencies, higher allele frequency differences between groups, and higher levels of heterozygosity compared to non-PGx variants genome-wide (Figure 2A-C). We considered group-specific differences in PGx variation in terms of the fixation index (*F_ST_*), a commonly employed measure of population differentiation, and effect allele frequency differences. PGx *F_ST_* and effect allele frequency difference values are highly correlated, as can be expected, and the largest differences are seen for the Black-White and Black-Hispanic group comparisons (Figure 2D-F). Notably, even the most extreme values of *F_ST_* fall well below 0.5, indicating the most PGx variation is found within rather than between SIRE groups. Nevertheless, there are 82 PGx variants that show statistically significant (FDR *q*<0.05) values of allele frequency differentiation between any individual SIRE group and the other two groups, *i.e.* their complements (Figure 2G). All-against-all pairwise distances for HRS participants were calculated using PGx variants and projected into two-dimensions with multi-dimensional scaling (MDS). *K-* means clustering was used to create three groups based on the PGx MDS distances, and individuals were labeled according to their SIRE (Figure 2H). Genome-wide patterns of PGx variation characterized in this way show 96.1% correspondence to SIRE group labels (Figure 2I).

### SIRE versus GA for Partitioning Pharmacogenomic Variation

Given the overall correspondence, and group-specific differences, seen for SIRE and GA, we wanted to compare the utility of SIRE versus GA for partitioning pharmacogenomic variation in the US. Here, we asked two questions regarding PGx variation between groups: (1) are PGx allele frequencies correlated between SIRE and GA groups, and (2) do GA groups partition PGx variation more so than SIRE groups? The first question was addressed by regressing PGx frequency differences between grouping systems (SIRE vs. GA groups), and the second question was addressed by considering the deviation of the regression from the unity line (*i.e.* the expected value under perfect correlation). As expected given the observed similarities between SIRE and GA groups, PGx allele frequency differences are highly correlated when corresponding group pairs are compared (Figure 3). The highest correlation is seen when the Black and White SIRE groups are compared to their corresponding GA groups. Comparisons that include the Hispanic SIRE group show lower levels of correlation.

**Figure 3.**
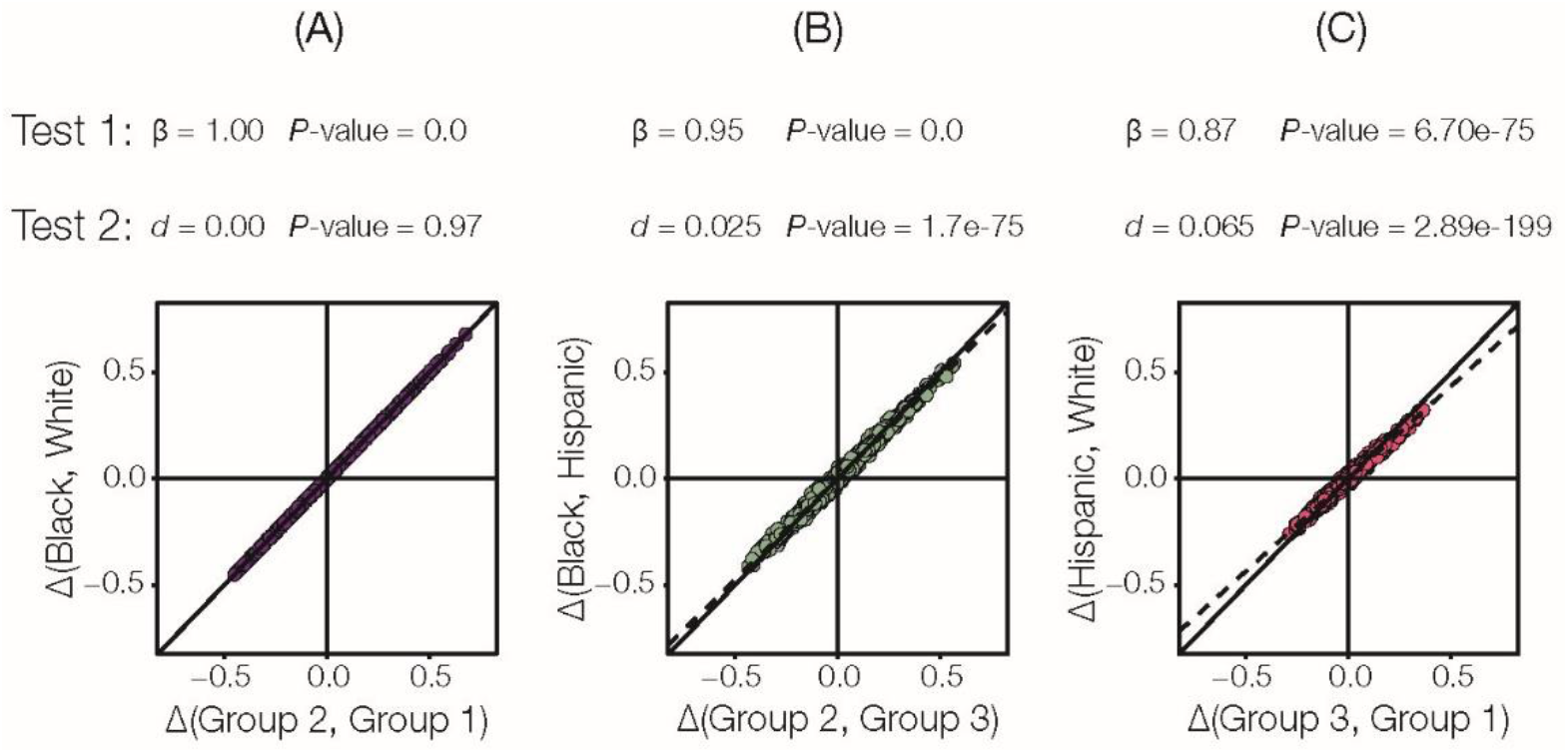
Self-identified race/ethnicity (SIRE) versus genetic ancestry (GA) for partitioning pharmacogenomic (PGx) variation. (A-C) Regression of pairwise PGx variant effect allele frequency differences calculated using SIRE (y-axis) versus the corresponding GA groups (x-axis). Results of two statistical tests are shown for each of three pairwise group regressions. Test 1 evaluates whether SIRE and GA PGx allele frequencies are correlated, and test 2 evaluates that amount of additional resolution on PGx variant divergence that is provided by GA compared to SIRE. Details on each test are provided in the text.

With respect to the second question regarding the partitioning of PGx variation, allele frequency differences between the Black/White SIRE groups and their corresponding GA groups fall almost entirely along the unity line; in this case, genetic ancestry does not provide any additional information regarding PGx variation (Figure 3A). For both comparisons that include the Hispanic group however, the slope of the regression is less than one, indicating greater PGx allele frequency differences between GA groups compared to their corresponding SIRE groups (Figure 3B and 3C). Thus, GA does provide more information than SIRE when ethnicity is considered, but the effect size of this difference is small (*d*=2.5% for Black/Group 2 vs. Hispanic/Group 3 and *d*=6.5% for Hispanic/Group 3 vs. White/Group 1).

Thus far, we have shown that SIRE and GA groups are highly concordant for the HRS cohort and that PGx allele frequency differences are similar for both classification systems. Since SIRE labels are routinely collected as patient provided information, and are also readily available as part of electronic health records, we focused on PGx variation between SIRE groups to explore the potential clinical utility of race and ethnicity. We wanted to know whether PGx effect allele frequency differences of the magnitude observed here have any utility for guiding medication prescription decisions in light of the fact that the majority of PGx variation is found within rather than between SIRE groups. We considered the odds ratios for the apportionment of PGx risk alleles among individual SIRE groups and their complements as an indicator of SIRE groups’ predictive utility, given that odds ratios are widely used to associate categorical risk factors with health outcomes^43^. We also computed absolute risk increase values to account for the population frequency of PGx risk alleles when considering the magnitude of between group differences as well as the accuracy with which SIRE group membership predicts PGx alleles or genotypes. Detailed results for all PGx variants analyzed here can be found in Supplementary Table 2.

Examples of highly differentiated PGx variants are shown in Table 2 and Figure 4. The relative percentages of PGx effect (above) and non-effect (below) alleles across SIRE groups reveal the extent of differentiation for these variants (Figure 4A), and the observed allele frequency differences are associated with SIRE groupspecific continental ancestry fractions (Figure 4B-D). Nevertheless, as described above and shown in Figure 2, even highly differentiated PGx variants show levels of *F_ST_* that indicate substantially more within than between group variation (see pie charts in Figure 4B-D). Despite the relatively high levels of within group PGx variation, these variants show high group-specific odds ratios and substantial absolute risk increase values. In other words, HRS cohort members’ racial and ethnic self-identities carry substantial information that can be used to stratify pharmacogenomic risk at the population level. However, the accuracy levels with which group affiliations predict specific risk alleles or genotypes are only marginally high, indicating that SIRE has relatively less utility for individual-level risk prediction compared to risk stratification.

**Figure 4.**
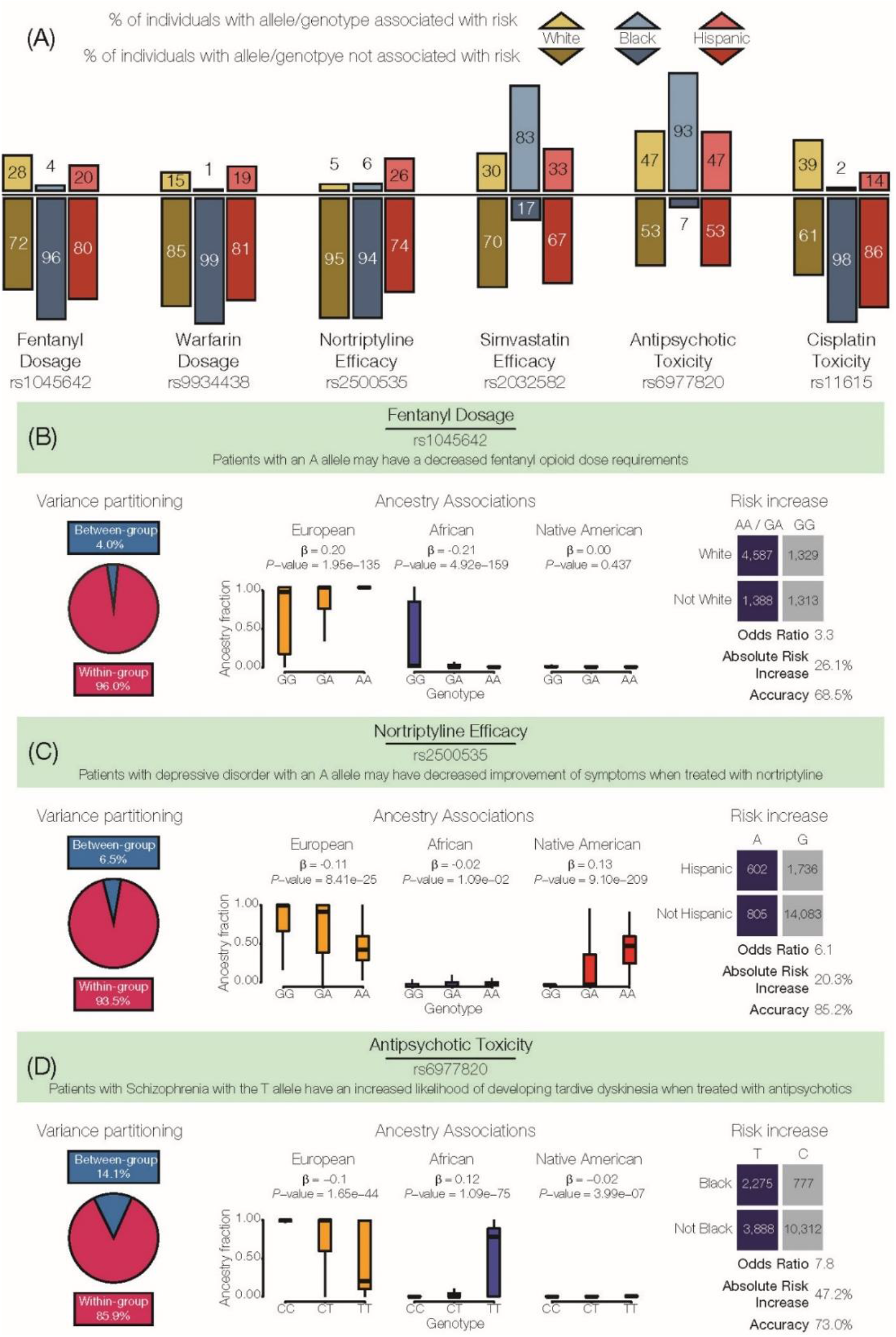
Examples of highly differentiated pharmacogenomic (PGx) variants. (A) SIRE group percentages of effect (above axis) versus non-effect (below axis) alleles/genotypes are shown for six highly differentiated PGx variants. Allele counts are used for the additive PGx effect mode, and genotype counts are used for the dominant effect mode. (B-C) The extent of within versus between group variation, ancestry associations, and PGx stratification/risk by SIRE groups are shown for three examples. Ancestry associations relate the ancestry fractions for individuals that bear distinct PGx genotypes: European (orange), African (blue), and Native American (red). Effect (blue) versus non-effect (gray) allele/genotype counts are compared for the group enriched for a specific PGx variant compared to the other two groups. Allele counts are shown for the additive PGx effect mode, and genotype counts are shown for the dominant mode. Group-specific allele/genotype counts were used to compute odds ratios and absolute risk increase values (risk stratification) along with group-specific prediction accuracy values (risk prediction) as shown

**Table 2.** Examples of highly differentiated PGx variants. This table lists some examples of highly diverged PGx variants in the three SIRE groups under consideration. In the table, ‘Ref. Pop.’ refers to Reference Population, OR refers to Odds Ratios, ARI

| rsID | Drug | Effect | Effect allele frequency |  |  | Ref. Pop. | OR | ARI | Accuracy |
| --- | --- | --- | --- | --- | --- | --- | --- | --- | --- |
|  |  |  | White | Black | Hispanic |  |  |  |  |
| rs1045642 | Fentanyl | Dosage | 0.78 | 0.37 | 0.70 | White | 3.26<br>(2.96, 3.60) | 26.1<br>(24, 28) | 68.5<br>(67.0, 69.9) |
| rs9934438 | Warfarin | Dosage | 0.38 | 0.83 | 0.33 | Black | 8.27<br>(7.18, 9.54) | 45.93<br>(44, 48) | 66.53<br>(65.03, 68.03) |
| rs2884737 | Warfarin | Dosage | 0.27 | 0.04 | 0.18 | Black | 8.99<br>(7.43, 10.87) | 36.0<br>(34, 38) | 52.5<br>(50.5, 54.5) |
| rs4646450 | Tacrolimus | Metabolism | 0.16 | 0.84 | 0.33 | Black | 66.80<br>(49.17, 90.88) | 63.15<br>(62, 65) | 71.5<br>(70.2, 7.2) |
| rs2500535 | Nortriptyline | Efficacy | 0.05 | 0.06 | 0.26 | Hispanic | 6.1<br>(5.40, 6.82) | 20.3<br>(18, 22) | 85.2<br>(84.6, 85.9) |
| rs11615 | Platinum compounds | Efficacy | 0.37 | 0.88 | 0.64 | Black | 9.90<br>(8.85, 11.09) | 45.95<br>(45, 47) | 63.5<br>(62.4, 64.6) |
| rs20455 | Atorvastatin | Efficacy | 0.36 | 0.79 | 0.40 | Black | 14.2<br>(11.11, 18.17) | 35.71<br>(34, 37) | 50.01<br>(47.9, 52.1) |
| rs1048943 | Capecitabine, Docetaxel | Efficacy | 0.04 | 0.02 | 0.27 | Hispanic | 12.74<br>(11.14, 14.79) | 39.4<br>(37, 42) | 87.3<br>(86.5, 88.1) |
| rs6977820 | Antipsychotics | Toxicity | 0.04 | 0.28 | 0.05 | Black | 14.8<br>(12.13, 18.14) | 45.96<br>(44, 48) | 60.09<br>(58.4, 6.1) |
| rs1801394 | Methotrexate | Toxicity | 0.46 | 0.72 | 0.67 | White | 2.82<br>(2.63, 3.02) | 24.68<br>(23, 26) | 59.40<br>(58.2, 60.1) |
| rs16969968 | Nicotine | Toxicity | 0.66 | 0.95 | 0.80 | Black | 8.17<br>(6.97, 9.59) | 26.6<br>(26, 28) | 43.17<br>(41.4, 44.9) |

refers to the Absolute Risk Increase percentage. Values in brackets specify the 95% confidence intervals for each computation.

For example, the A allele of the PGx variant (rs1045642) in the ATP Binding Cassette Subfamily B Member 1 (ABCB1) encoding gene is associated with a decreased fentanyl opioid dose requirement^47^ (Figure 4B). This PGx variant has a dominant mode of effect, such that patients with either the AA or GA genotype tend to metabolize fentanyl slower than patients with the GG genotype and will therefore require a lower dosage. 96.0% of variation for this PGx variant is partitioned within SIRE groups compared to 4.0% variation between groups. However, the dosage-associated genotypes are far more common in individuals who identify as White (*OR*=3.3, *CI*=3.0-3.6; *ARI*=26.1%, *CI*=24.0%-28.3%), and from the ancestry association plot, it can be seen that the effect allele (A) is highly correlated with European genetic ancestry (β=0.20, *P*=1.95e-35). Self identification as White predicts dosage-associated genotypes with 68.5% accuracy.

Similarly, a PGx variant (rs2500535) in the Uronyl 2-Sulphotransferase (*UST*) gene has been found to be associated with the efficacy of nortriptyline – an antidepressant – in patients with major depressive disorder^48^ (Figure 4C). This PGx variant has a dominant mode of effect; patients with the A allele are associated with a decreased improvement of depression symptoms when prescribed nortriptyline. These lower efficacy genotypes are more common in individuals who identify as Hispanic. Even though the variation at this genomic site is far higher within (93.5%) compared to between (6.5%) groups, the odds ratio for having risk-associated genotypes is high for the Hispanic population (*OR*=6.07, *CI*=5.44-6.82) along with a high absolute risk increase (*ARI*=20.3%, *CI*=18.5%-22.2%). Hispanic ethnicity predicts nortriptyline efficacy-associated genotypes with 85.2% accuracy.

Another PGx variant (rs6977820) found in the Dipeptidyl Peptidase Like 6 (*DPP*) gene has been associated with adverse response to antipsychotic drugs (Figure 4D). This PGx variant has an additive effect mode, whereby the T allele is positively correlated with African ancestry and associated with tardive dyskinesia among Schizophrenia patients treated with antipsychotics ^49^. When individuals that self-identify as Black are compared to the other two SIRE groups, most variation at this variant is found within (85.9%) rather than between (14.1%) groups. However, the odds ratio for the presence of the risk allele for adverse reaction to antipsychotics is high (*OR*=7.7, 95% *CI*=7.1-8.49), as is the absolute risk increase (*ARI*=47.2%, 95% *CI*=45.4%-48.9%), consistent with a substantially elevated risk of adverse drug reaction for the Black SIRE group compared to the others. Individuals who self-identify as Black can be predicted to have the effect-associated allele with 73.0% accuracy.

### Clinical Value of Pharmacogenomic Stratification by SIRE

We quantified the clinical utility of SIRE for partitioning PGx variation by comparing the ability to predict PGx effect alleles/genotypes before (pre) and after (post) stratification of the population by SIRE. The approach we used is equivalent to the comparison of pre- and post-test probabilities for diagnostic tests, where the test in this case is patient stratification by SIRE. For any given PGx variant, the pre-test probability is the overall population prevalence of the PGx effect allele/genotype, and the post-test probabilities are the group-specific positive predictive values (PPVs) for the PGx effect allele or genotype. Allele counts were used to compute these probabilities for PGx variants that show an additive effect mode, and genotype counts were used for the dominant effect mode. The absolute difference of the pre- and post-test probabilities calculated in this way was taken as a measure of the amount of information that is gained, with respect to PGx variant prediction for each specific group, when SIRE is used for patient stratification.

When highly differentiated PGx variants (Figure 2G and Figure 4) are analyzed in this way, the SIRE groups that show the highest effect allele frequencies for any given variant provide substantial additional information for PGx prediction. Considering the PGx variant (rs2500535) that is associated with Nortriptyline efficacy (Figure 4C), stratification by Hispanic identity yields an additional 14 individuals, for every 100 patients to be treated, who are predicted to show decreased improvement of symptoms related to depressive disorder. The information gain is even more extreme for the PGx variant (rs6977820) that is associated with antipsycohtic toxicity (Figure 4D). For this variant, stratification of individuals that self-identify as Black will yield an additional 39 out of every 100 patients that are counter-indicated for the antipsychotic medications owing to toxic side effects. The overall levels of information gained via stratification by SIRE differ widely by group. Individuals that self-identify as Black show the highest levels of information gain for PGx variant prediction followed the Hispanic and White groups, respectively (Figure 5). This pattern can be attributed to the relative numbers of individuals in each SIRE group together with the extent of genetic diversification seen between groups. The relatively high frequency of PGx effect alleles (Figure 2A) also contributes to the amount information gain observed here, given the fact that PPVs depend on the prevalence of the condition that is being tested (i.e. the presence of PGx effect alleles/genotypes).

**Figure 5.**
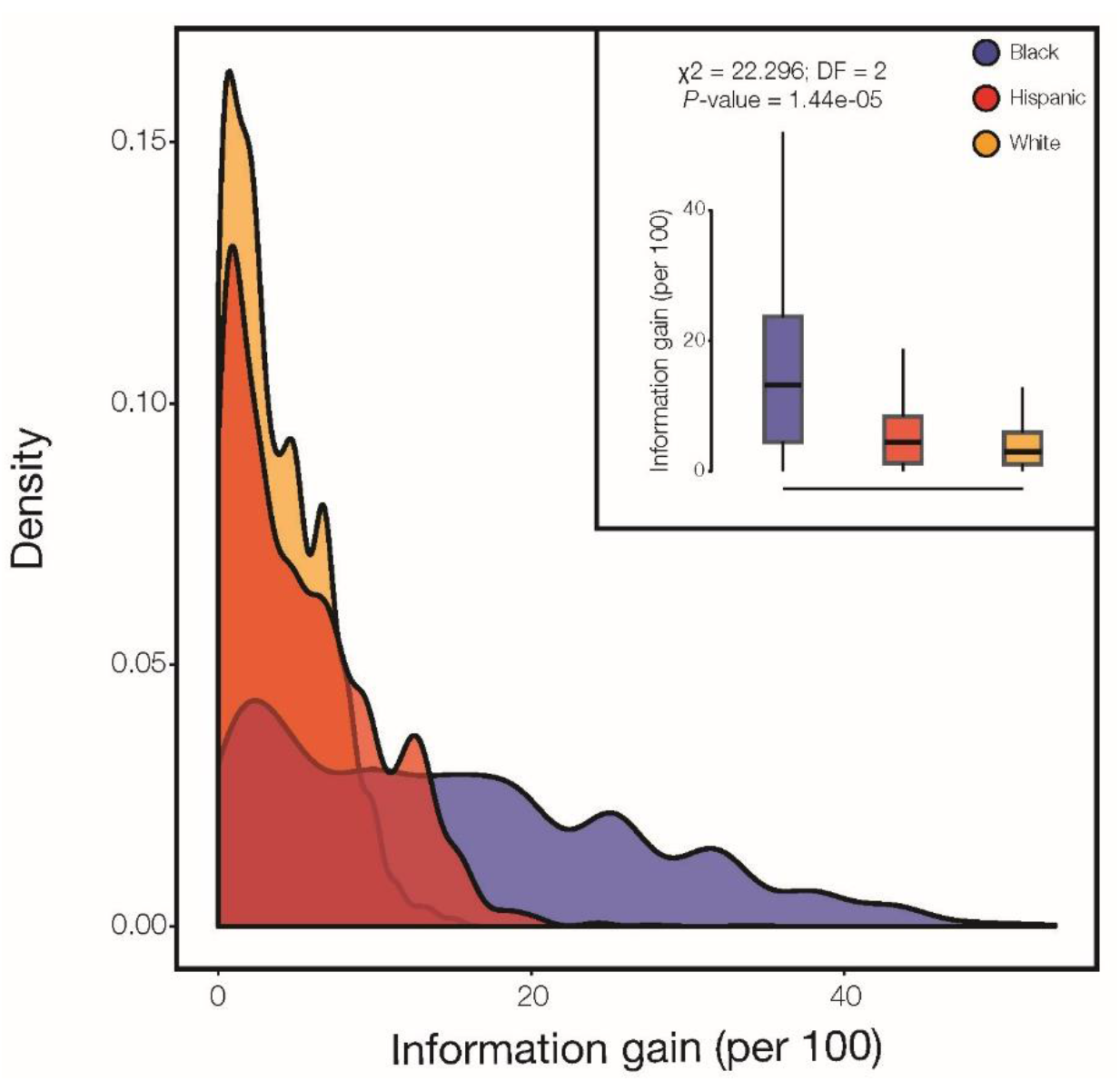
Information gained when SIRE is used for PGx stratification. The amount of information gained per 100 individuals is the number additional correct PGx variant predictions made when SIRE is used to stratify the population. Information gain is calculated for all PGx variants in each SIRE group, as described in the text, and the group-specific distributions are shown as density distributions and box-plots (inset): White (orange), Black (blue), and Hispanic (red).

## Discussion

### Concordance Between SIRE and GA in the US

The SIRE and GA groups from the US analyzed here show >96% overall concordance (Figure 1, Supplementary Figures 2 and 3). It must be stressed that these results only apply to the three major racial/ethnic groups covered by the ~8,600 individual HRS cohort; nevertheless, the concordance between SIRE and GA seen for the HRS cohort is very much consistent with a number of previous studies of the US population. In 2005, investigators showed a 99.9% concordance between SIRE and genetically derived clusters for 3,636 individuals from four racial/ethnic groups groups^50^, and a 2007 study reported 100% classification accuracy of individuals from geographically separated population groups when thousands of genetic variants were used for clustering^51^. More recently, a study of >11,000 cancer patients from The Cancer Genome Atlas found an 95.6% concordance between self-reported race (not ethnicity) and GA^52^, and a massive study of >200,000 individuals from the Million Veterans Program found >99.4% concordance between SIRE and GA^53^. The latter two studies relied on machine learning classifiers powered by vectors of 7 and 30 ancestry principal components, respectively, whereas our clustering algorithm uses vectors of only three continental ancestry components to classify individual genomes. Additionally, the distribution of GA fractions observed here for the HRS cohort SIRE groups is consistent with previous studies^34^;^46; 54^–^56^. Taken together, our results and others underscore the extent to which continental ancestry patterns can distinguish SIRE groups in the US.

Genetic differences accumulate among populations when they are reproductively isolated, and isolation by distance^57^ best accounts for the apportionment of human genetic diversity among global populations^58^. Populations that are physically distant, or separated by major geographic barriers, are more genetically diverged than nearby populations^59^. It follows that the appearance of population structure, *i.e.* distinct clusters of genetically related individuals, can represent an artifact of uneven sampling of human populations at extremes of distance^60^. For instance, isolation by distance can explain much of the apparent genetic structure observed for major genome sequencing projects such as the 1000 Genomes Project^35^;^61^ and the Human Genome Diversity Project^36; 62^. Conversely, when human populations are sampled more evenly across a range of distances, and in the absence of major geographical barriers, genetic diversity appears to be continuously distributed as a cline of variation^63^;^64^.

Isolation by distance can be taken to explain the concordance of the SIRE and GA groups observed for the HRS cohort, since the three major US SIRE groups are made up of individuals with ancestry from continental population groups – European, African, and Native American – that were isolated at great distances for tens-of-thousands of years before coming back together over the last 500 years^34; 46^. Since each SIRE group contains distinct patterns of continental ancestry, they correspond well to objectively defined clusters formed based on the partitioning of GA (Figure 1, Supplementary Figures 2 and 3). In addition, despite the fact that these population groups are currently co-located within the US, assortative mating based on culture stands as an ongoing reproductive barrier among groups^65^;^66^ (but see below for an important caveat regarding this fact). It is nevertheless important to note that most of the SIRE and GA groups analyzed here are not composed of individuals with highly coherent ancestry patterns. Only the White/Cluster 1 groups show coherent ancestry patterns, whereas the Black/Cluster 2 and Hispanic/Cluster 3 groups are made of up of individuals that vary along a range of continental ancestry fractions (Figure 1 and Supplementary Figure 2). This is especially true for the Hispanic group, consistent with the fact Hispanic is an intentionally broad label that covers individuals from different races and with very distinct ancestry patterns^67^.

An important caveat with respect to the high concordance between SIRE and GA observed here relates to the age of the individuals in the HRS cohort (Table 1). We chose to focus on older Americans given their disproportionate use of prescription medications^31^, and HRS recruited participants aged 50 and over starting in 1992. The average age of the HRS cohort analyzed here is 57.5 years (CI: 57.0-58.0), and all of the study participants were born before 1965, when there were still “anti-miscegenation” laws in nineteen states^68^. Rates of intermarriage among SIRE groups have increased substantially since that era^69^, and as admixture continues to increase over time, the ancestral coherence of SIRE groups is expected to fall precipitously. Increased rates of immigration, coupled with the arrival of more globally diverse immigrant groups, will also blur boundaries between SIRE groups, potentially rendering the current labels clinically uninformative. Indeed, the most widely used SIRE labels in the US are mandated by the OMB, and they will likely be revised in the near future to better capture the increasing diversity of the US population. As such, the clinical relevance of SIRE will almost certainly decrease over time.

### Within versus between group genetic divergence

It has long been appreciated that the vast majority of human genetic variation is found within rather than between populations. This fundamental result was first reported for worldwide racial groups, based on analysis of a handful of (surrogate) genetic markers^70^, and has since been confirmed by numerous studies of populations defined by GA using larger-scale analyses^62^;^71–75^. The distinction between this fundamental result and the high concordance seen for SIRE and GA, as well as the ability to cluster human population groups at various levels of relatedness, can be explained by the difference between univariate methods for variance partitioning versus multivariate classification methods^76^;^77^. The analysis of PGx variation reported here is univariate, since we focus on the apportionment of variation for individual PGx variants, and we confirm that the majority of PGx variation is found within the HRS cohort groups (Figure 2 and 4).

We used a standard evidence based medicine analytical framework^43^;^44^ in an effort to understand the clinical relevance of PGx variation that is partitioned among SIRE groups in this way. In particular, we asked how the observed PGx differences between groups could be clinically relevant when the majority of variation falls within population groups, even for the most divergent variants found here. Despite the observed pattern of within versus between group PGx variation, we found numerous cases of high odds ratios and high absolute risk increases for the group-specific prevalence of PGx variants (Table 2 and Figure 4). In other words, membership in any given SIRE group can entail substantially greater odds, and far higher risk, of carrying clinically relevant PGx variants compared to members of other groups. Information of this kind should be an important consideration for clinicians charged with making treatment decisions and could also be of value for well-informed patients.

Finally, it should be emphasized that humans are far more similar than they are different at the genomic level, both within and between population groups. As of August 2019, there were 674 million annotated single nucleotide variants among the ~3 billion sites in the human genome^78^. Thus, more than 75% of genomic positions are conserved among all human population groups, and for those positions that do vary, the majority are rare variants that segregate at <1% frequency worldwide^35^. Nevertheless, the results reported here underscore the potential clinical relevance for the small the minority of genetic variants that show relatively high levels of between-group divergence.

### Caveats and limitations

It is important to note that in this study we measure the frequency of PGx variants across different SIRE and GA groups, rather than drug response differences *per se.* Even though the penetrance of PGx variants is generally high^2^, clinical interpretations of variant frequency differences should be considered in light of variable penetrance levels as well. In cases of low penetrance, the magnitude of drug response differences between groups will be dampened. Furthermore, if PGx variants have different magnitudes of effect for different groups, *i.e.* group-specific effect sizes, then differences in drug response cannot be directly inferred from PGx variant frequency differences alone. However, since the majority of PGx variants are causative protein coding variants^2^, the likelihood of group-specific effect sizes is far lower than would be expected for non-coding variants discovered by genome-wide association studies, which are typically tag markers that are linked to nearby causative variants. Finally, the focus on single nucleotide variants (SNVs) is another limitation of the study, given the fact structural variants and multi-variant haplotypes have also been associated with inter-individual drug response differences. Nevertheless, the vast majority of PGx variants annotated in the PharmGKB database are SNVs^2^, suggesting that our analytical approach captures most of the known variant-drug associations.

### The current and future utility of race and ethnicity in pharmacogenomics

As previously noted, demographic trends in the US suggest that the clinical relevance of SIRE, including its predictive utility for PGx variation, is expected to continuously decrease over time. The increasing adoption of routine genetic testing for precision medicine could also render SIRE obsolete for stratifying PGx variation^79^. This is because genotyping of specific PGx variants will obviously provide far more accurate risk prediction than SIRE. For example, even a highly divergent PGx variant, like the antipsychotic toxicity associated variant rs6977820 (Figure 4D), will yield a mis-prediction of the PGx risk allele 27% of the time if SIRE alone were used as a predictor. In this sense, the high group-specific PGx odds ratios and absolute risk increases observed in this study are best considered as surrogate guides to inform the optimal choice of prescribed medication, rather than precise diagnostic tools. In other words, SIRE categories provide valuable information for stratifying PGx risk at the population level but not for predicting individual-level PGx variants. Having said that, and despite the promise of population scale genomic screening initiatives and biobanks^80^, such as the NIH All of Us project^81^, the day when all Americans will have ready access to their genetic profiles remains far in the future. Unfortunately, this is likely to be even more so for minority communities that are vastly underrepresented among clinical genetic cohorts^82^;^83^. Until that time, SIRE will remain an important feature for clinicians to consider when making treatment decisions.

Perhaps most importantly, the current utility of SIRE is most apparent for groups who are underrepresented in biomedical research. Individuals who self-identify as Black or Hispanic stand to gain far more information with respect to precision treatment decisions than those who identify as White (Figure 5). This finding can be attributed to the relative frequencies of individuals in each of the three SIRE groups analyzed here, which closely mirror the current demography of the US, and the extent of genetic divergence among groups. If a ‘one size fits all’ approach to drug prescription is used, patients who identify as White are more likely to receive the most appropriate treatment, since their PGx variant frequencies will be closest to the overall population mean. Conversely, individuals who identify as Black or Hispanic have the most to lose if SIRE is not considered when making treatment decisions.

## Supplemental Data

Supplemental data included two tables and four figures.

## Declaration of Interests

The authors declare no competing interests

## Acknoweldgements

The authors thank Dr. Greg Gibson for comments on a manuscript draft and Dr. Joe Lachance for helful discussion.

---

**Supplementary Table S1.**
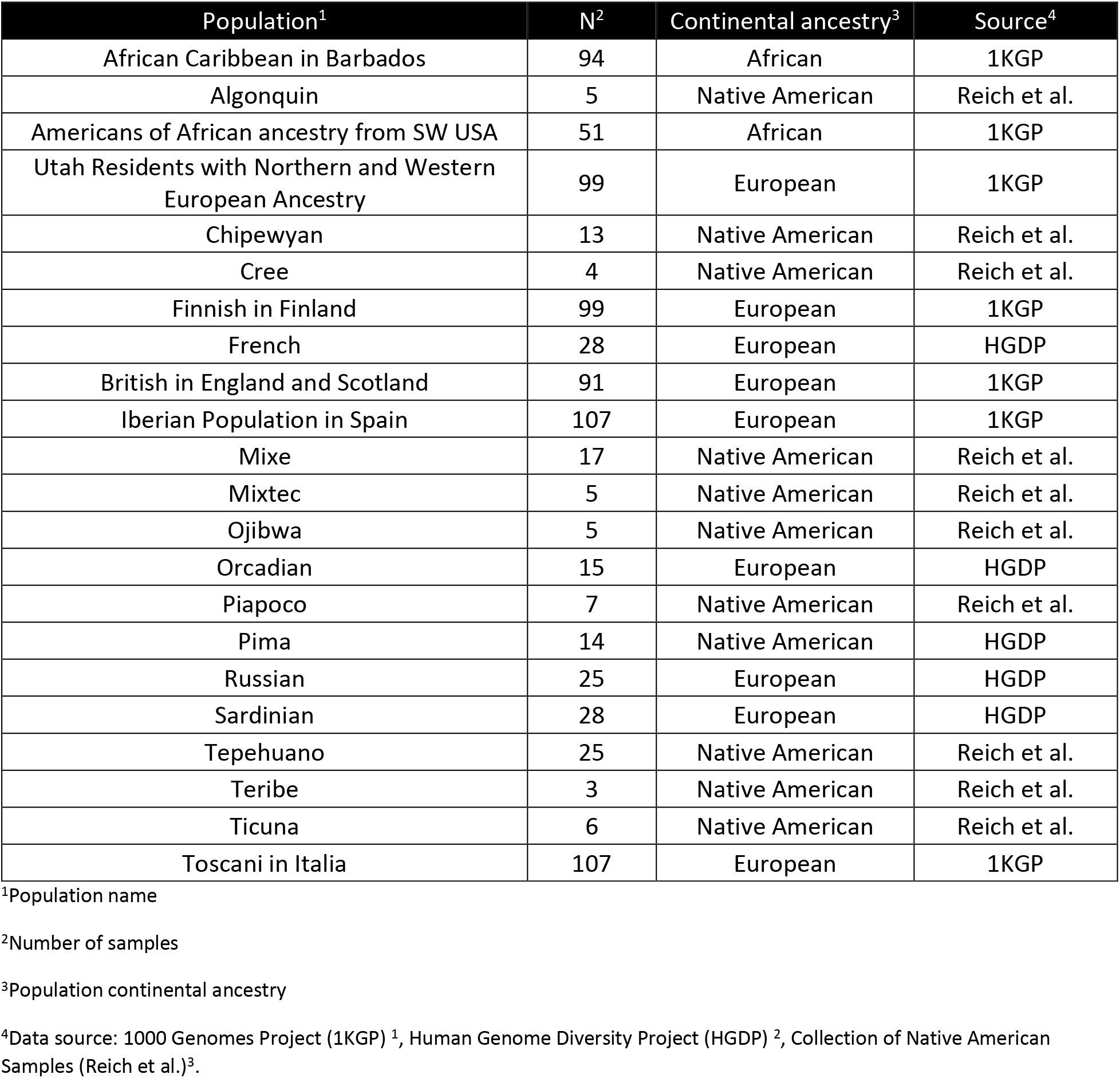
Global reference populations used for genetic ancestry inference.

| Population<sup>1</sup> | N<sup>2</sup> | Continental ancestry<sup>3</sup> | Source<sup>4</sup> |
| --- | --- | --- | --- |
| African Caribbean in Barbados | 94 | African | 1KGP |
| Algonquin | 5 | Native American | Reich et al. |
| Americans of African ancestry from SW USA | 51 | African | 1KGP |
| Utah Residents with Northern and Western European Ancestry | 99 | European | 1KGP |
| Chipewyan | 13 | Native American | Reich et al. |
| Cree | 4 | Native American | Reich et al. |
| Finnish in Finland | 99 | European | 1KGP |
| French | 28 | European | HGDP |
| British in England and Scotland | 91 | European | 1KGP |
| Iberian Population in Spain | 107 | European | 1KGP |
| Mixe | 17 | Native American | Reich et al. |
| Mixtec | 5 | Native American | Reich et al. |
| Ojibwa | 5 | Native American | Reich et al. |
| Orcadian | 15 | European | HGDP |
| Piapoco | 7 | Native American | Reich et al. |
| Pima | 14 | Native American | HGDP |
| Russian | 25 | European | HGDP |
| Sardinian | 28 | European | HGDP |
| Tepehuano | 25 | Native American | Reich et al. |
| Teribe | 3 | Native American | Reich et al. |
| Ticuna | 6 | Native American | Reich et al. |
| Toscani in Italia | 107 | European | 1KGP |

<sup>1</sup>Population name
<sup>2</sup>Number of samples
<sup>3</sup>Population continental ancestry
<sup>4</sup>Data source: 1000 Genomes Project (1KGP) <sup>1</sup>, Human Genome Diversity Project (HGDP) <sup>2</sup>, Collection of Native American Samples (Reich et al.)<sup>3</sup>.

**Supplementary Figure 1.**
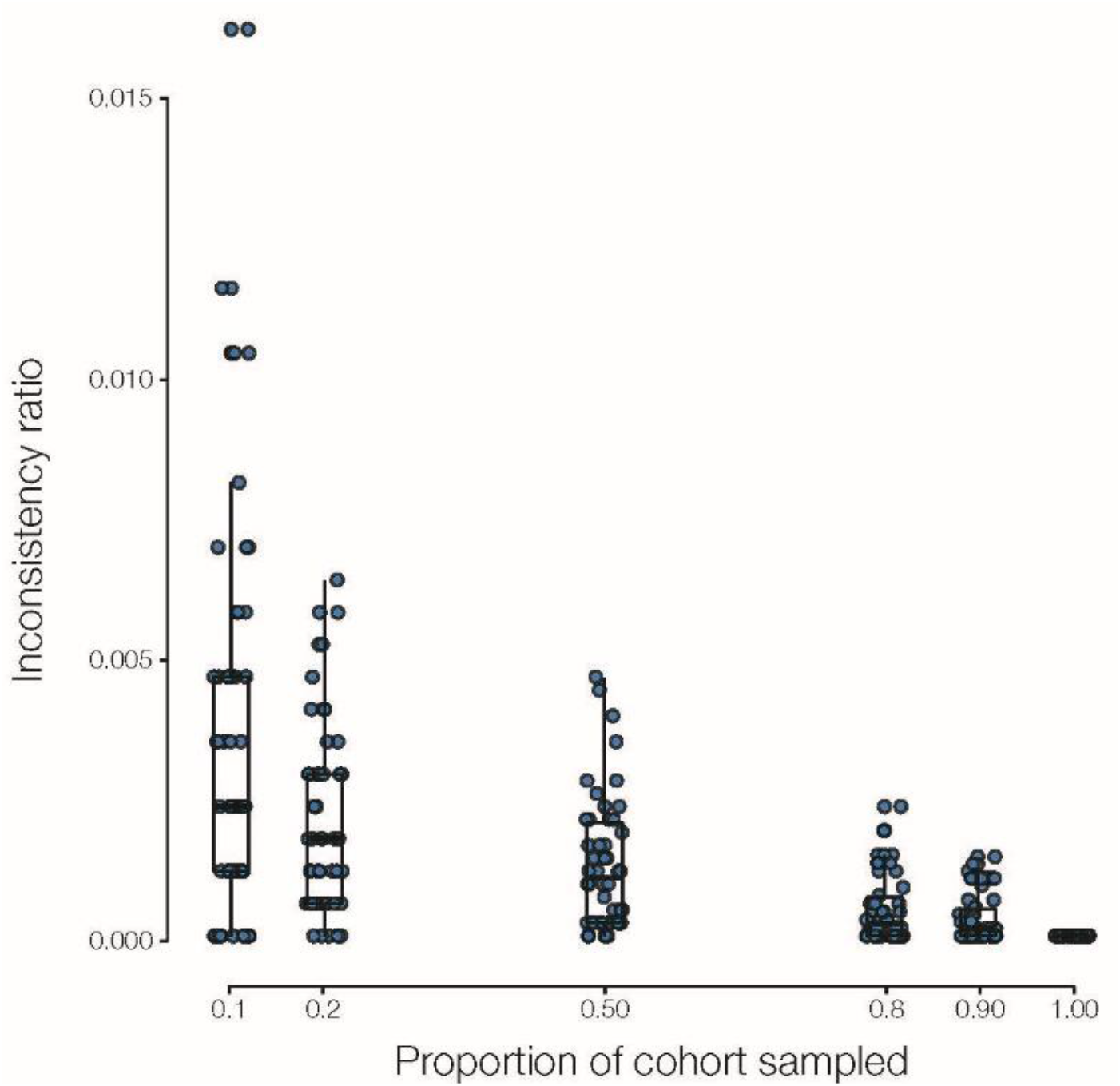
Permutation analysis to evaluate the stability of *k*-means genetic ancestry (GA) clusters. The HRS cohort was randomly sampled at different proportions, where the proportion of the cohort sampled = the number of participants in the random sample / the total number of participants in the cohort. For each random sample, *k*-means clustering was run 50 times and an inconsistency ratio was calculated for each independent run, where the inconsistency ratio is the number of mismatches between the random sample group assignments / the number of participants in the random sample. In other words, the inconsistency ratio measures the error in *k*-means cluster assignments due to sampling bias. As can be expected, error is higher for smaller random cohort proportions and decreases monotonically as the proportion of the random cohorts increases. Nevertheless, the error level, even at the smallest sampling proportions, is extremely low. The mean error at a sampling proportion of 0.1 is 0.4%, and when the entire cohort is sampled (i.e. cohort proportion=1) *k*-means clustering is 100% consistent.

**Supplementary Figure 2.**
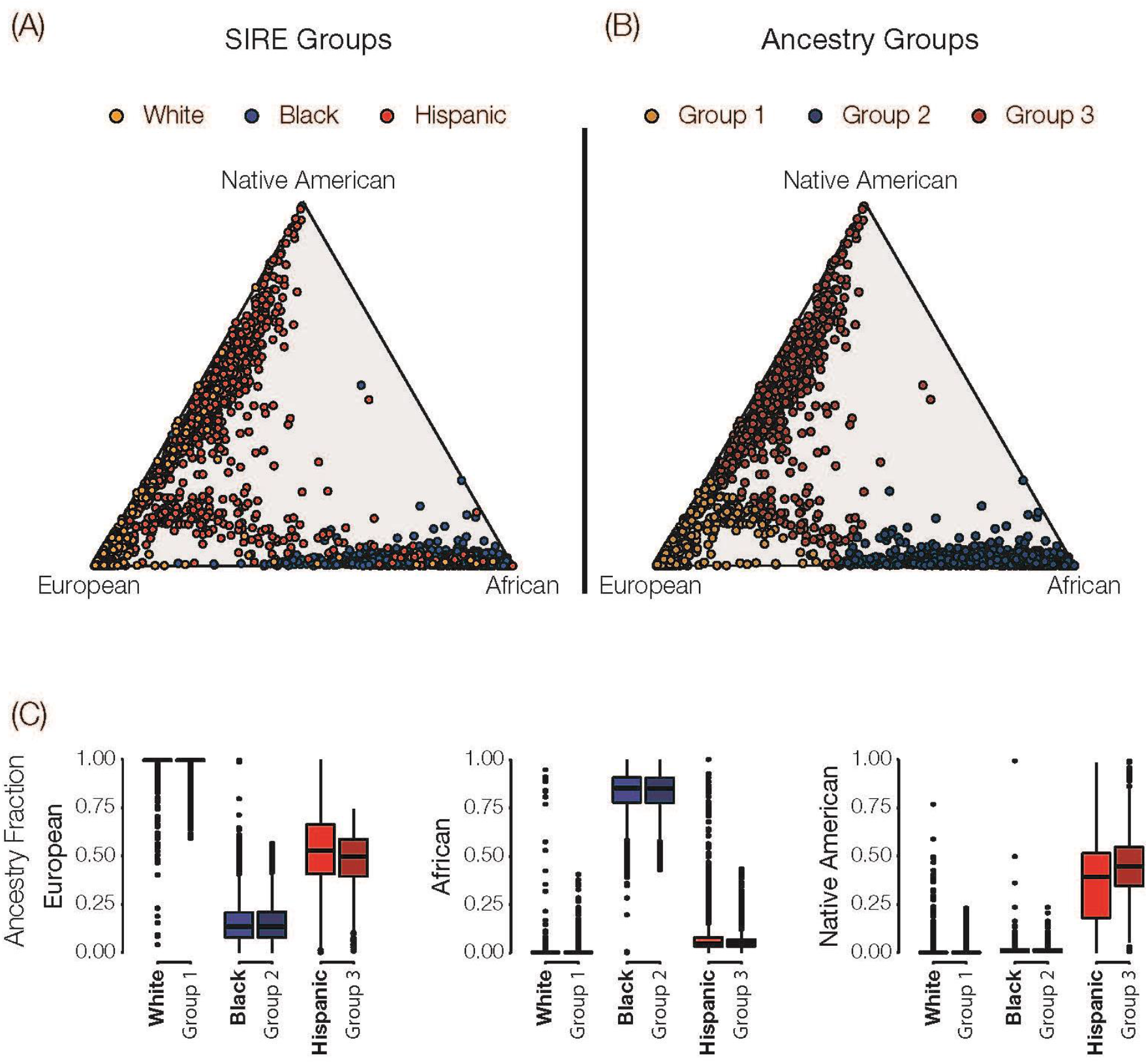
Comparison of self-identified race/ethnicity (SIRE) versus genetic ancestry (GA) groups in the US. Ternary plots showing the relative continental ancestry fractions for HRS participants are shown with individuals color coded by SIRE (A) or genetic ancestry (B). SIRE and their corresponding GA groups are coded as White/Group 1 (orange), Black/Group 2 (blue), and Hispanic/Group 3 (red). (C) Distributions of continental ancestry fractions - European, African, and Native American - for HRS participants are shown corresponding SIRE and GA groups.

**Supplementary Figure 3.**
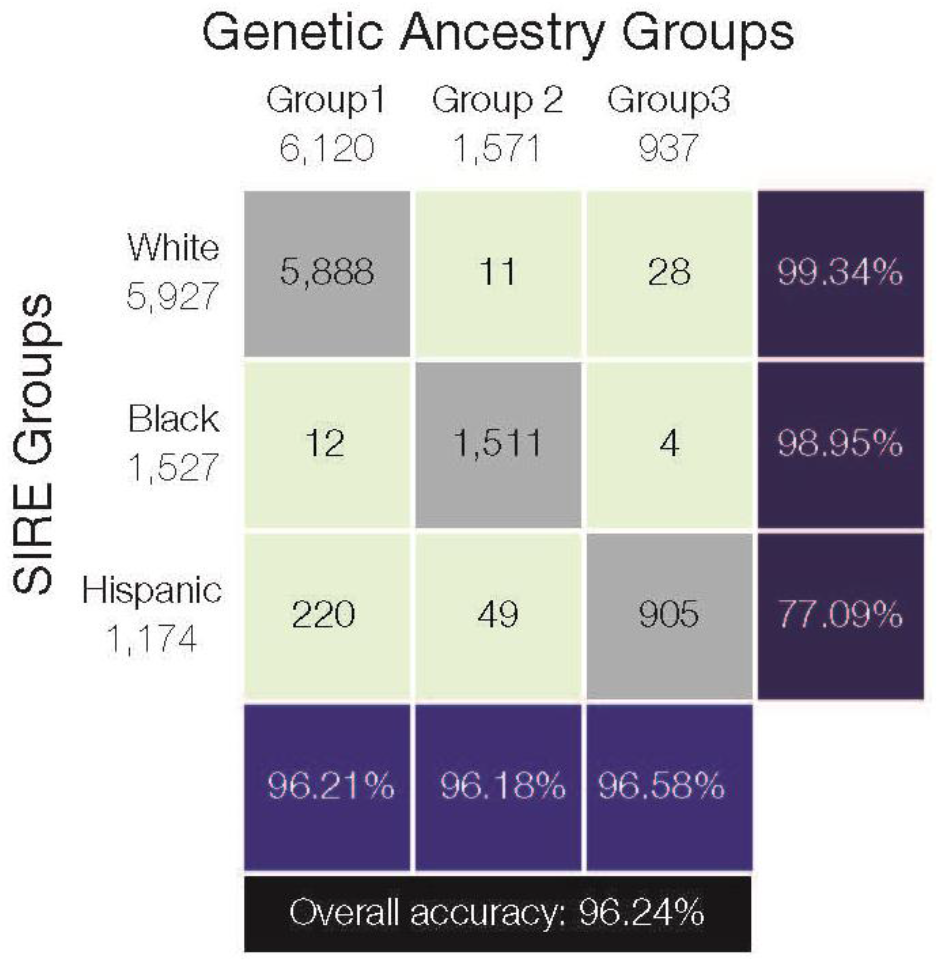
Correspondence between self-identified race/ethnicity (SIRE) versus genetic ancestry (GA) groups in the US. Numbers of HRS participants that fall into each combination of SIRE and GA groups is shown along with the percentage correspondence. Individual percent correspondence values are calculated as the number of individuals along the diagonal, i.e. that fall into the corresponding SIRE and GA groups, divided by the total number of individuals in each SIRE group (right) or each GA group (bottom), times 100. The overall percent correspondence is calculated as the number of individuals along the diagonal divided by the total number of individuals in the HRS cohort, times 100.

**Supplementary Figure 4.**
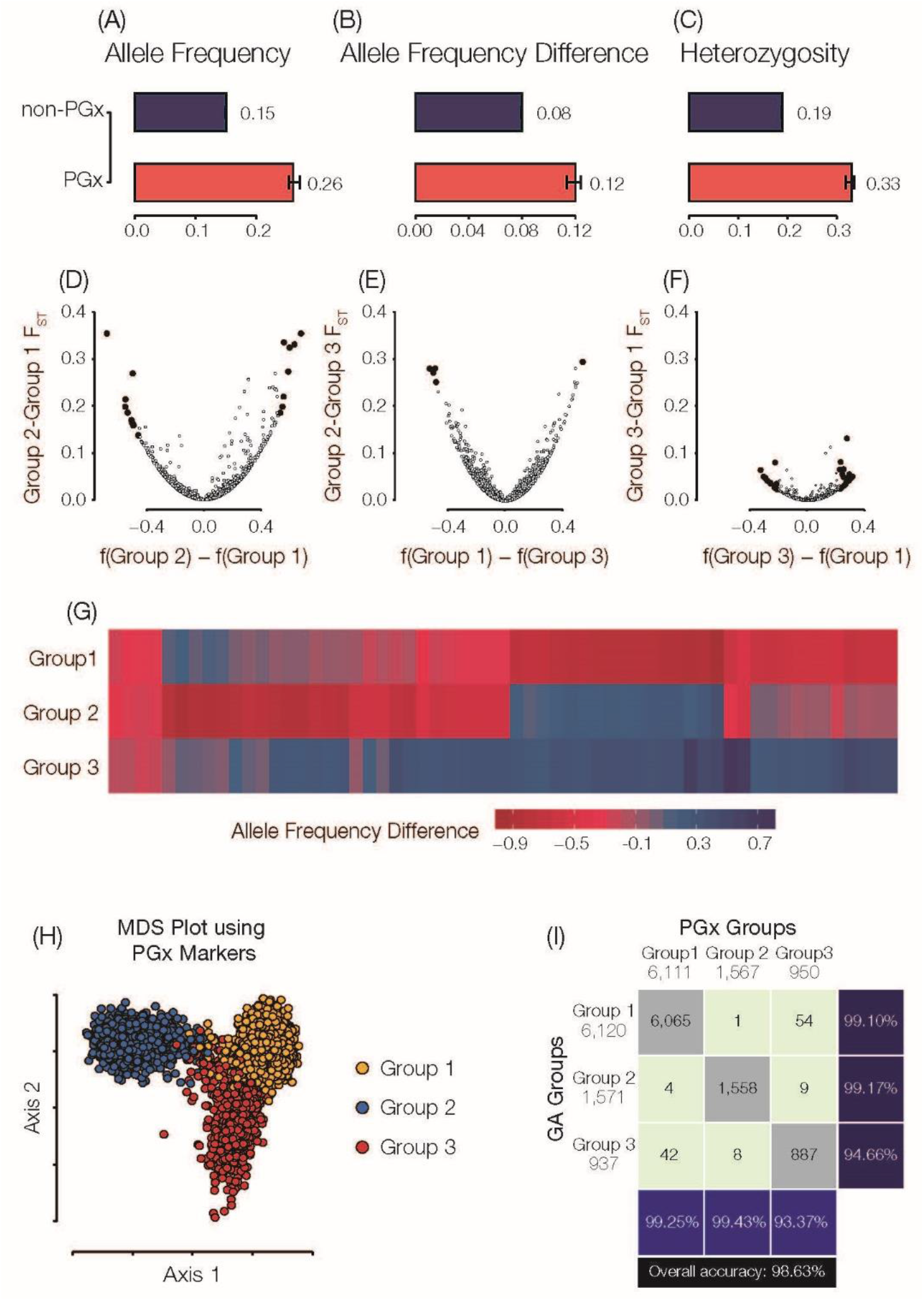
Pharmacogenomic variation in the US: genetic ancestry (GA). Data shown here correspond to GA groups; analogous results for SIRE groups shown in Figure 2. Genome-wide average allele frequencies (A), group-specific allele frequency differences (B), and heterozygosity fractions (C) are shown for PGx variants (red) compared to non-PGx variants (blue). (D-F) Fixation index (F_ST_; y-axis) and allele frequency differences (x-axis) for pairs of GA groups. Statistically significant PGx allele frequency differences are highlighted in black. (G) Heatmap showing group-specific allele frequencies for significantly diverged PGx variants. (H) Multi-dimensional scaling (MDS) plot showing the relationship among individual genomes as measured by PGx variants alone. Each dot is an individual HRS participant genome, and genomes are color-coded by participants GA groups. (I) The correspondence between GA groups and PGx groups defined by K-means clustering on the results of the MDS analysis.

